# The Markov link method: a nonparametric approach to combine observations from multiple experiments

**DOI:** 10.1101/457283

**Authors:** Jackson Loper, Trygve Bakken, Uygar Sumbul, Gabe Murphy, Hongkui Zeng, David Blei, Liam Paninski

## Abstract

This paper studies *measurement linkage*. An example from cell biology helps explain the problem: imagine for a given cell we can either sequence the cell’s RNA or we can examine its morphology, but not both. Given a cell’s morphology, what do we expect to see in its RNA? Given a cell’s RNA, what do we expect in its morphology? More broadly, given a measurement of one type, can we predict measurements of the other type? This measurement linkage problem arises in many scientific and technological fields. To solve this problem, we develop a nonparametric approach we dub the “Markov link method” (MLM). The MLM makes a conditional independence assumption that holds in many multi-measurement contexts and provides a way to estimate the *link*, the conditional probability of one type of measurement given the other. We derive conditions under which the MLM estimator is consistent and we use simulated data to show that it provides accurate measures of uncertainty. We evaluate the MLM on real data generated by a pair of single-cell RNA sequencing techniques. The MLM characterizes the link between them and helps connect the two notions of cell type derived from each technique. Further, the MLM reveals that some aspects of the link cannot be determined from the available data, and suggests new experiments that would allow for better estimates.

**Significance Statement:** Novel experimental techniques are developing quickly, and each technique gives new perspectives. Ideally we would build theories that account for many perspectives at once. This is not easy. One challenge is that many experiments use measurement techniques that alter or destroy the subject, making it impossible to measure the same subject with both techniques and difficult to combine data from different experiments. In this paper we develop the Markov Link Method, a new tool that overcomes this challenge.

## Introduction

Many scientific experiments produce data that come from different measurement techniques. To use such data, we need to understand the link between the different types of measurements. For example, in a biological context, a transcriptomic-morphological link would answer a question like: ‘We have measured the transcriptome of this cell - what might its morphology have been?’

This paper formalizes the idea of a measurement link with a conditional probability. Suppose we obtain a measurement *x* from one technique. Given this measurement, *g(y | x)* is the conditional probability of obtaining *y* from the second technique, when applied to measure the same specimen. The problem we solve is how to estimate this conditional distribution.

It is particularly challenging to estimate the link when multiple measurements of the same specimen are unavailable. For example, it is currently often difficult to observe both the RNA expression and morphological properties of a single cell. Rather, we typically have two datasets: one contains the transcriptomes of one group of cells and the other contains the morphology of a another group of cells from the same population. Traditional statistical tools offer no way to use these two distinct datasets to estimate the transcriptomic-morphological link. Figure 1 sketches this difficult situation, which is common in biological problems [1]. This general *measurement linkage* problem arises in many scientific and technological fields where it is difficult to take multiple types of measurements of the same specimen.

**Figure 1:**
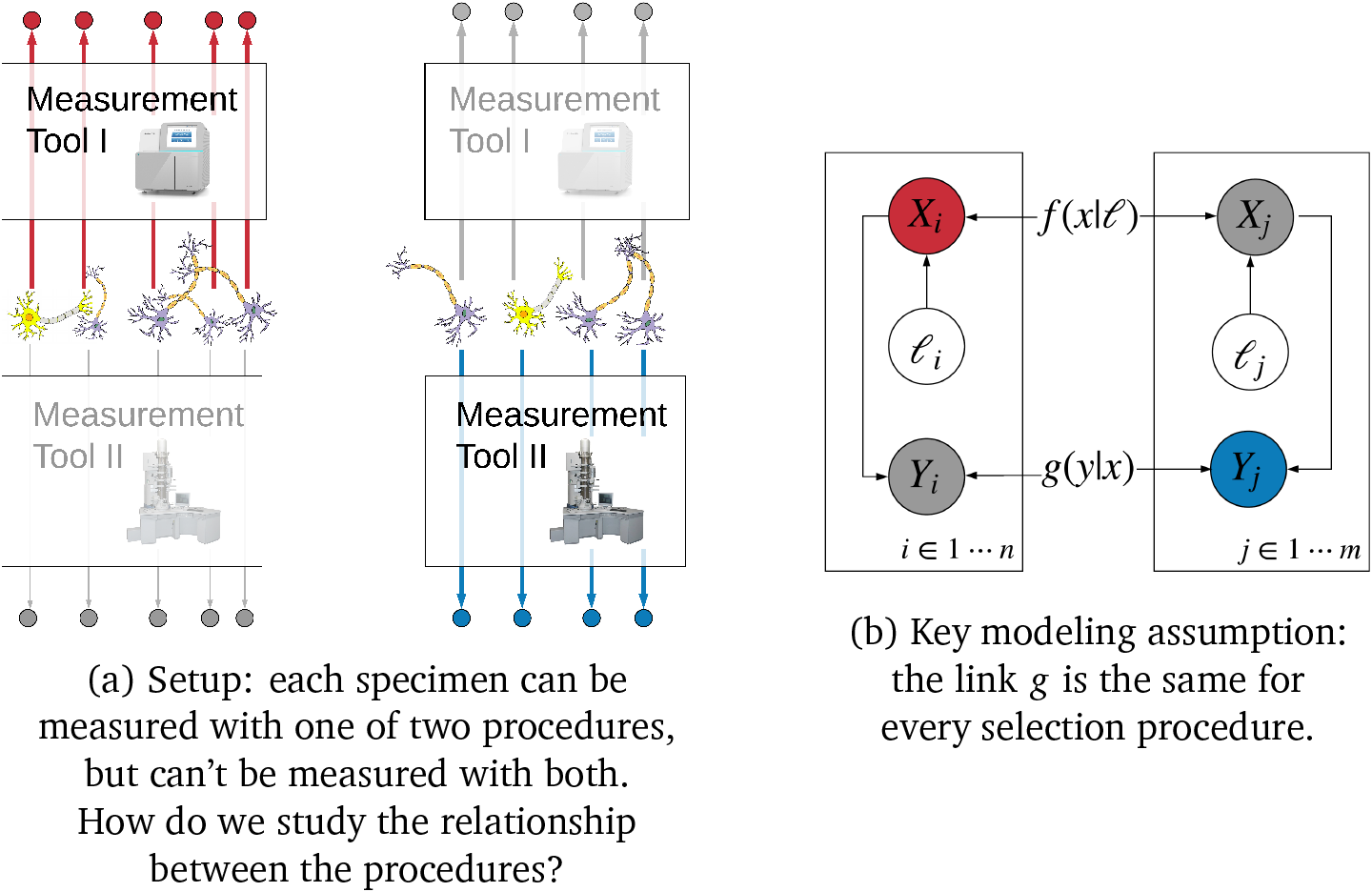
The Markov link method (MLM) links different measurement procedures. In subfigure (a) we have a population of neurons. We select half of the neurons and determine what type of neuron they are using transcriptomic measurements. We classify the second half of neurons based on their morphological properties. The MLM provides a way to link these two types of measurement and reconcile the two resulting notions of cell-type. In subfigure (b) we present a graphical model diagram articulating the statistical assumption upon which our analysis is based (cf. [2] for a thorough explanation of this type of diagram). For each neuron in each group, this diagram considers the selection procedure (ℓ), the transcriptomic type (*X*), and the morphological type (*Y*). The assumption is that the transcriptomic type is sufficiently detailed to *statistically isolate* the transcriptomic type from the selection procedure. This is reflected in the arrows of the diagram; all paths from ℓ variables to *Y* variables pass through *X* variables. The assumption allows us to estimate the link, *g*.

To address this problem, we develop the Markov link method (MLM). The MLM estimates the link nonparametrically. It relies only on a natural conditional independence assumption that can be expected to hold in a wide variety of scientific contexts.

Here is how it works. Statistically, the challenge is that we never observe samples from the full joint distribution of the two types of measurements. But suppose we can condition on an external variable ℓ and then sample from the conditional distributions of each type, given ℓ. Now the key idea behind the MLM is that we can exploit a natural Markov assumption – that the link itself is unaffected by the external variable ℓ – to mathematically relate the link to the conditional distributions of each type. Therefore, when this Markov assumption holds, we can estimate the link without ever observing samples from the full joint distribution.

In the next section, we describe the MLM in more detail and discuss examples where its assumptions are plausible. Then we develop statistical theory, establishing conditions under which the MLM estimator is statistically consistent (i.e., estimates the correct link given sufficient data), and show that the estimator provides accurate measures of uncertainty. Finally, we apply the MLM to real data generated by a pair of single-cell RNA measurement procedures. The results help characterize the link between them, and suggest new experiments that further improve the estimated link.

## 1 The Markov link method

The Markov link method uses a collection of experiments to estimate the link between two measurement procedures. As a running example, consider biological experiments studying single cells. Assume each experiment can be characterized by:

- A *selection procedure* defines how we select cells to measure. For example, one experiment selects liver cells, another selects cells which express the Vip protein, and another selects cells that exhibits a particular proteomic marker.
- A *measurement procedure* defines what we measure about each cell. One procedure measures the transcriptomic profile of a cell and another measures the cell morphology.

Assume that for each selection procedure exactly two experiments have been conducted: one measures transcriptomics and the other measures morphology. There are no experiments where an individual specimen was measured with both measurement procedures. This setup is sketched in Figure 1. For each cell gathered in any experiment, we are interested in:

1. ℓ, the selection procedure used to obtain the cell.
2. *X*, the cell’s ‘transcriptomic type,’ the result of applying a transcriptomic measurement procedure to the cell (e.g. single-cell RNA sequencing), followed by an assignment of the cell to one of a finite number of cell types defined in terms of transcriptomic properties.
3. *Y*, the cell’s ‘morphological type,’ the result of applying a morphological measurement procedure to the cell (e.g., tracing the cell and counting the number of branches and segments), followed by an assignment to a cell type defined in terms of morphological properties.

The task is to learn the link *g* that connects *X* and *Y*. The challenge is that for each cell we can only observe *either X* or *Y*, but not both. The MLM overcomes this challenge to link the transcriptomic and morphological perspectives on ‘cell-type.’ With the MLM, we can make statements of the form ‘This cell has transcriptomic type 3; therefore it probably has morphological type F.’

We begin with a critical assumption, what we call the *Markov link assumption* we assume that the link between measurement procedures is unaffected by the choice of selection procedure. In mathematical language, we assume there exists some *g*(*y|x*) so that for every selection method ℓ,

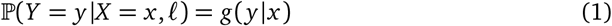

where 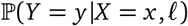 indicates the probability that the morphological type is *y* given that the transcriptomic type is *x* and the specimen was selected with method ℓ. The graphical model in Figure 1 embodies the assumption. In the running example, it says that once we know the transcriptomic type *x* of a given cell, learning the procedure ℓ that selected the cell provides no additional information about its morphological type *y*. (Section 1.1 discusses further examples where this conditional independence assumption is reasonable.)

The Markov link assumption is useful because it yields equations that connect what we are interested in (i.e., *g*) with the data we have. This data is governed by the following conditional distributions:

- *f*(*x*|ℓ) - the probability that a cell sampled with procedure ℓ will have transcriptomic type *x*;
- *h(y*|ℓ) - the probability that a cell sampled with procedure ℓ will have morphological type *y*.

Under the Markov link assumption, these distributions are connected to *g* by the equations

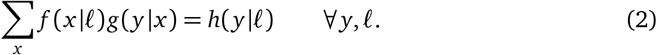

If we knew *f* and *h* then we could (under suitable assumptions) invert these equations to estimate the link. Let’s denote this (conceptual) estimator as *ĝ_f,g_*, to emphasize the fact that it depends upon *f, h*.

In practice we do not know *f* and *h*. We take a Bayesian perspective and treat these as unobserved variables. For prior beliefs, we assume a noninformative uniform prior 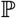 and we then incorporate new knowledge by conditioning on the observed knowledge. We have two important pieces of knowledge about *f* and *h*. First, we have observed the data, which we denote as 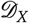 and 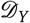. Second, a consequence of the Markov link assumption is that there exists *some* value *g* such that Equation (2) holds; let 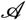 denote the event that this holds. By conditioning on this knowledge, we obtain the posterior distribution 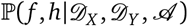. Now we can define a point estimate for the link by computing, e.g., the mean of this posterior:

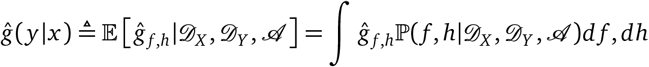

We can use similar strategies to estimate credible intervals *C_x,y_* for the link.

In summary, for each transcriptomic type *x* and morphological type *y*, the Markov link method uses the experimental data to produce two objects:

1. *ĝ*(*y|x*) - a point estimate for the link *g*(*y|x*);
2. *C_x,y_* - a credible interval which contains the true link *g*(*y |x*) with high probability. A provides the full algorithm for estimating these quantities.^1^

### 1.1 Other examples

Our running example has been the problem of linking two ways of measuring cell-types. However, the MLM is a general approach for synthesizing experimental data where the Markov link assumption is valid. To illustrate these broader applications, we discuss some other examples where the MLM could potentially be applied:

- Quality control for manufacturing. Consider a manufacturing process that produces components; to evaluate production quality, they use two different destructive tests. The first kind of test is cheap, but its accuracy is unknown. A more established test is accurate but expensive. The manufacturer is interested in the link: given results from the cheap test (*X*), what might have been the results from the accurate test (*Y*)? To estimate the link, the MLM uses different batches of production ℓ. For example, ℓ might indicate the day on which a component was made. The assumption: the link doesn’t depend on the day the component was made. This is reasonable if the tests are performed on different days. In this case, the MLM can be used to calibrate the two kinds of tests.
- Astronomy photography. Consider an astronomer taking photographs of stars over time; depending on availability, some pictures are taken using one camera and other pictures are taken using a different camera. To standardize the photos, the photographer is interested in the link: given a photo from one camera (*X*), what might the picture have looked like if it was taken with the other camera (*Y*)? To estimate the link, the MLM uses different astronomical targets ℓ. For example, ℓ might indicate which star is being photographed. The assumption: the difference between the cameras is the same for each star. This is reasonable as long as the camera settings are set to the same values for each star. In this case, the MLM can be used to standardize photographs from different cameras.
- Personalized medicine. Consider two sources of knowledge about how transcriptomes might help us personalize cancer medication. We can measure the transcriptome of cancer cells from inside humans, but experimenting on humans is difficult and morally fraught. We can also culture immortalized cells from human cancers and measure their transcriptomes; experimenting on immortalized cells is cheap and straightforward. To generalize work on immortalized cells to human patients, we need the link: given a cultured cell’s transcriptome (*X*), what can we expect about the corresponding in-vivo transcriptome (*Y*)? To estimate the link, the MLM can use different types of cancer (ℓ). The assumption: the transcriptomic effects of culturing the cells is the same for each cancer type. This maybe reasonable as long as the cancer types are sufficiently similar. In this case, the MLM can be used to help generalize results from immortalized cells to real human patients.
- Replication crisis and lab effects. We would like to understand the differences in how two labs perform experiments. These differences can be understood by looking at the link: given results from one lab (*X*), what results could we expect from another lab (*Y*)? To estimate the link, the MLM can use different specimen batches (ℓ). For example, ℓ might indicate a batch of mice, half of which were sent to one lab and half of which were sent to another lab. The assumption: the lab effects are the same for each batch of mice. In this case, the MLM can be used to discover “lab effects” that impede successful experimental replication.

Note that the marginal distribution of *Y* may be different for each selection procedure. The MLM assumption only requires that the *conditional* distribution of *Y*|*X* is the same. In the language of statistics: the measurement *X* is ‘sufficient’ for the measurement *Y*.

## 2 Mathematical Results and Simulations

We would like the Markov link method estimators to have three properties:

**Estimator convergence**: As we obtain more samples, *ĝ*(*y|x*) should converge to the true link *g*(*y|x*) for each *x, y*.

**Interval concentration**: As we obtain more samples, the credible interval *C_x,y_* should get smaller. Asymptotically it should include nothing but the point *g*(*y|x*).

**Conservative coverage**: *g*(*y|x*) ∈ *C_X_*,*_y_* with high probability.

In the next sections we explore these properties, theoretically and empirically.

### 2.1 Theoretical results

Conservative coverage is the most essential of the three properties. It ensures that that the method doesn’t ‘lie.’ The method returns an interval *C_x,y_* which is supposed to contain the true value of the link parameter *g*(*y|x*). Conservative coverage ensures that this is indeed the case, with high probability.

The MLM achieves asymptotically conservative coverage under two mild conditions: *positive probability* (*g*(*y*|*x*) > 0 for every *x*,*y*) and *linear independence* (the vectors *{f*(·|ℓ)}ℓ are linearly independent).^2^ By asymptotically conservative, we mean that as the number of samples goes to infinity any ε-inflation of *C_x,y_* is guaranteed to include the truth with arbitrarily high probability. A statement and proof of our conservative coverage theorem can be found in Appendix B.

What about the other two properties, estimator convergence and interval concentration? In many cases, the MLM achieves both of these conditions. However, in other cases, we can show that an issue known as ‘non-identifiability’ blocks even the possibility of estimator convergence or interval concentration. This problem arises when there are not enough distinct selection procedures. Each additional selection procedure gives us a collection of bounds on the link *g*. If we do not have enough selection procedures, it may be impossible to recover the link exactly. We demonstrate these issues in detail in Appendix C.

To understand how the MLM may work in practice, we supplement these theoretical results with two empirical simulations designed to show the strengths and weaknesses of the method. We discuss these simulations in the next two sections.

### 2.2 Simulation I: Number of samples and number of selection procedures

We suppose we have two measurement procedures which return categorical measurements among six categories. The first procedure yields *X* ∈ {1,2,3,4,5,6} and the second yields *Y* ∈ {1,2,3,4,5,6}. To see how the method performs in different circumstances, we run many trials. In each trial we fix the number of selection procedures and pick a ‘ground truth’ link uniformly at random from the parameter space. We then simulate a dataset from this ground truth link, fixing the total number of samples (spreading these samples equally among all combinations of selection procedure and measurement procedure). Finally, we apply the MLM to the simulated data to get the point estimates *ĝ*(*y|x*) and the credible intervals *C_x,y_*. We measure the overall estimator convergence using a type of total variation distance:

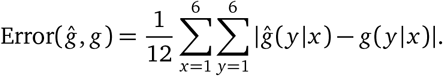

This error ranges between zero and one: zero indicates that *ĝ* = *g*, one indicates that the estimate has completely incorrect beliefs about the probability mass, i.e. *g*(*y|x*) = 0 whenever *ĝ*(*y|x*) > 0 and vice-versa. We also study at the MLM credible intervals and see how often they cover the true parameters.

Figure 2 summarizes the results. In trials with more samples, the estimator usually has lower error. However, only with at least six distinct selection procedures does the error converge to zero. This figure also shows that the credible intervals work correctly regardless of the number of samples or selection procedures; they include the ground truth with high probability. With many samples and selection procedures, the intervals are small and concentrated around the truth. With fewer samples or selection procedures, the intervals are typically larger. However, the uncertainty is not the same for every aspect of the link. In some cases we obtain a tight credible interval for *ĝ*(*y|x*) for some values of *x, y* and very loose intervals for other values.

**Figure 2:**
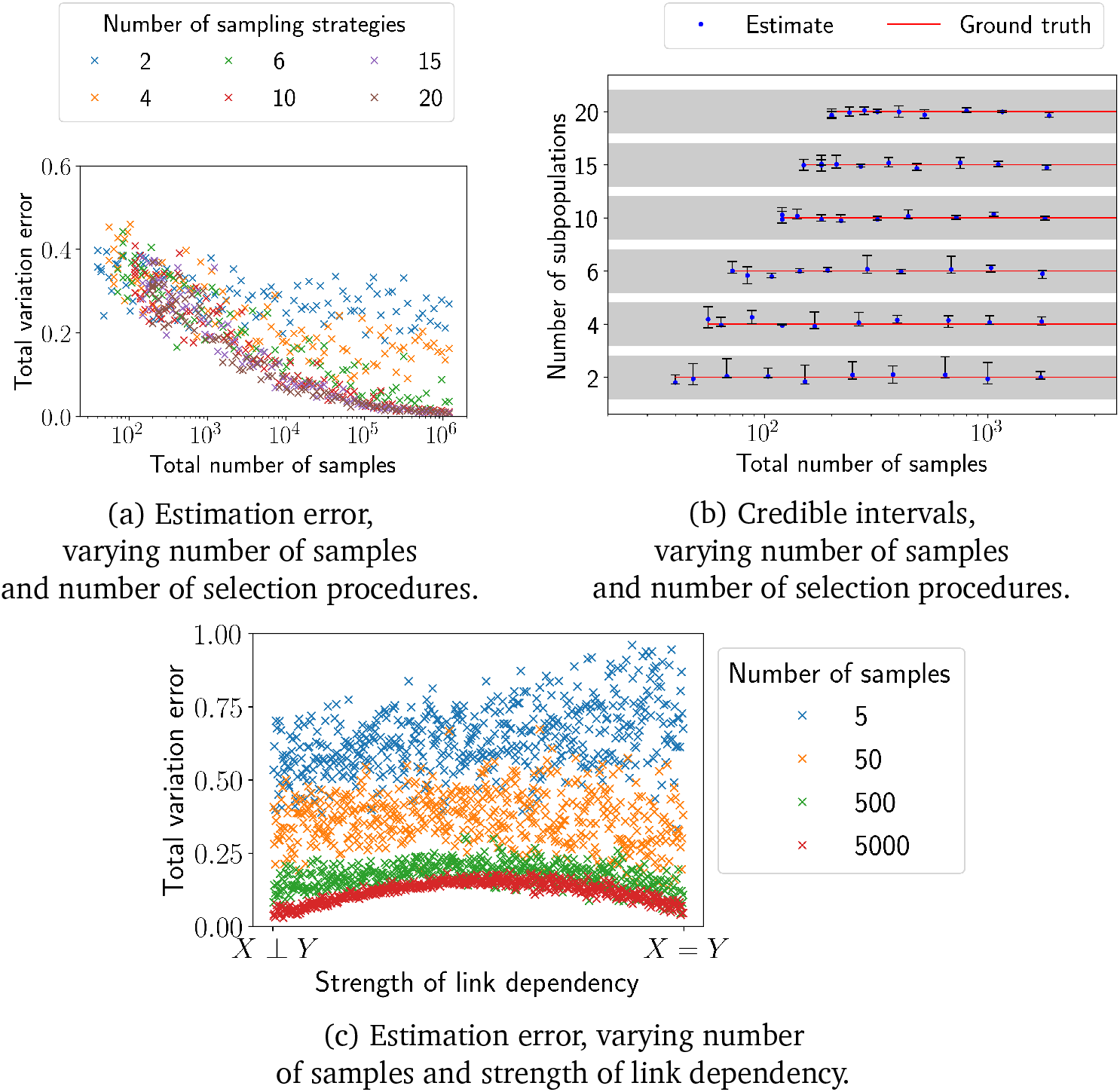
When can the MLM accurately estimate the link? We use simulations to test the MLM. In subfigure (a) we look at the error of the MLM link estimator in many different situations, varying the numbers of samples, selection procedures, and the link itself. In each trial the ‘ground truth’ link is picked uniformly at random from the parameter space. The MLM estimator accurately recovers the ground truth from simulated data, as long as we have enough samples *and* enough distinct selection procedures. In subfigure (b) we consider the MLM’s credible intervals. For a selection of trials we choose a random value of *x, y* and then compare the true link *g(y | x)* with the MLM interval *C_x,y_*. We show the interval (in black) centered around the ground truth (in red). Since the link probabilities lie in [0,1], the interval is always contained within ±1 of the ground truth (this larger region is shown in gray). The intervals mostly contain the truth, and get narrower with more samples as long as there are enough distinct selection procedures. Note that the horizontal axis indicates *total* number of samples. We only run the procedure when there are at least 20 samples per subpopulation; for this reason there are no intervals shown for small numbers of samples with many subpopulations. In subfigure (c) we focus on the case of exactly 4 selection procedures. In this case it is only possible to get very accurate estimates if the link takes on a particular form. If the link is independent (i.e. *X, Y* are independent) or if the link is invertible and deterministic (e.g. *X* = *Y*) then we can determine what the link is. However, in more challenging intermediate cases, small numbers of selection procedures may make it impossible to determine the true link; this issue is discussed in detail in Section C.

### 2.3 Simulation II: Different kinds of ground-truth links

The convergence of the estimator depends upon the link itself. Each trial of this simulation uses four selection procedures and the measurement procedures yield one of six categories. In the previous simulations we saw that estimator convergence was impossible in some cases, due to the small number of selection procedures - but those simulations picked the link uniformly from its parameter space. Now we will be more choosy.

On one extreme, we will produce trials where the link makes *X* and *Y* independent, i.e. *g(y|x)* = *g*(*y|x’* for every *x, x’*. On the other extreme, we will have trials where *X* and *Y* are deterministically related by the equation *X* = *Y*. We will also consider every link ‘in-between’ these two extremes (found by convex combinations). Figure 2 shows that estimator convergence is possible in the two extreme cases. However, estimator convergence fails for the in-between cases. In these cases there does not exist any consistent estimator, due to non-identifiability issues discussed in Appendix C.

We further examine one interesting case: let the measurements return one of 2^*k*^ categories and let us use only *k* +1 carefully chosen selection procedures. Now suppose the link defines any invertible deterministic function between *X* and *Y*. In this case, with enough samples, we can determine both that the relationship is deterministic and the exact specification of the invertible function. This result is proven in Appendix C, Theorem 2.

## 3 Empirical results for cell-types

Every cell in a human body has the same DNA (to a first approximation), but some cells behave differently from others. The biomolecular processes that drive this diversity are an area of active research [4, 5, 6]. Efforts such as the Human Cell Atlas project seek to map out a taxonomy of cell-types [7], thereby enabling a more systematic study of cellular diversity. Single cell transcriptomics provide an essential tool for reasoning about cell-types [8]. However, there are many different ways to measure transcriptomics, and most existing approaches destroy cells in the process of measuring their transcriptomic data. This makes it difficult to understand how taxonomies defined by one method may be related to taxonomies arising from another method. The Markov Link Method provides a new way to quantitatively approach these issues.

We will examine two procedures in particular, which we’ll call ‘Standard’ and ‘Patch.’ See [3] for details of these two methods. Briefly, the ‘Standard’ cell-typing pipeline applies single-cell

RNA sequencing to a population of cells and then applies clustering methods to divide the cells into types. The Patch pipeline is based on the ‘Patch-seq’ approach in [9]; these methods can obtain transcriptomic, electrophysiological, and morphological properties at the single-cell level, but the richness of the data comes at the cost of a somewhat degraded transcriptomic signal, leading to somewhat coarser cell-type determinations.

Both Standard and Patch methods produce cell-type determinations, but how can we check if these two methods produce consistent results? For example, Patch has a notion of a ‘Lamp5’ cell type. Standard gives a more granular analysis, dividing this type into many sub-types, such as ‘Lamp5 Pdlim5’ and ‘Lamp5 Slc35d3.’ If a cell was designated as ‘Lamp5 Pdlim5’ using Standard, we would hope that it be given the ‘Lamp5’ type by Patch. Unfortunately, since we cannot apply both methods to the same cell, we cannot directly test this question. The MLM gives a way to proceed, as long as we can use both methods on cells gathered with a variety of selection procedures.

We applied the MLM to a dataset that included both Standard and Patch data. Specifically, [10] describes a method for selecting different subpopulations of neurons. Each selection procedure yielded groups of cells with different proportions of the different cell-types. For each selection procedure and each measurement procedure, a number of cells were collected and typed using either the Standard or Patch pipeline. The result of this process was two tables, subsets of which are shown in Figure 3.

**Figure 3:**
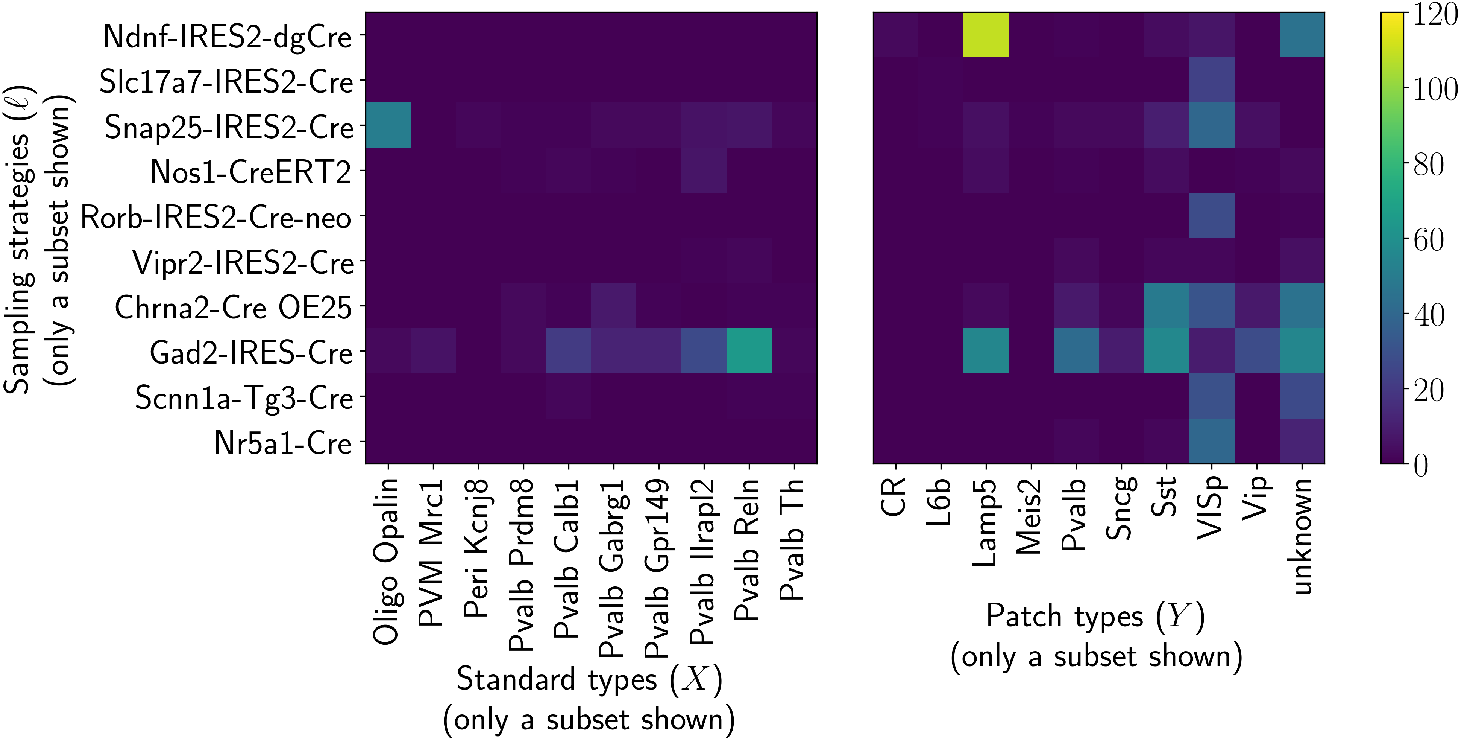
The input to the Markov link method: one experiment for each measurement procedure and each selection procedure. Here we show a portion of the real data to which we applied the MLM. Neurons from the visual cortex of mice were harvested using a variety of Cre/Lox-based selection procedures (cf. [3]). Each strategy was designed to sample from different subpopulations of cells. Neurons were measured to determine their ‘type,’ using one of two procedures: ‘Standard’ or ‘Patch.’ ‘Standard’ outputs 104 different types of neurons. ‘Patch’ has a coarser notion of cell-type, distinguishing only 10 types. For each experiment, we tabulated the number of cells assigned to each type. Above we show a subset of these results; the color of each square indicates the number of specimens found to have a particular type. Using this kind of data, the task is to calibrate the two classification protocols. That is, we want to be able to ask question of the following form: ‘if a neuron is classified as being of type ‘Peri Kcnj8’ by Standard, how might it have been classified by Patch?’

Given this data, we estimated the link between the Patch and Standard cell-typing pipelines. We calculated both a point estimate *ĝ*(*y|x*) and credible intervals *C_x,y_*. In Figure 4 we visualize these objects for selected values of *x, y*. For the Standard type *x* =‘Vip Rspo4’ and the Patch type *y* =‘Vip,’ we have that *C_x,y_* = [.88,1.0]. This supports the idea that the true link satisfies *g*(*y|x*) ≥ .88: at least 88% of the cells classified as type ‘Vip Rspo4’ by the Standard method will be classified as ‘Vip’ by the Patch method. However, for other types there is more ambiguity. For the Standard type *x* =‘Lamp5 Egln 1’ and the Patch types *y* =‘Lamp’ we have *C_x,y_* = [0,1.0]. The data do not give a definitive answer as to whether cells with Standard type ‘Lamp5 Egln 1’ are being classified with Patch type ‘Lamp.’

**Figure 4:**
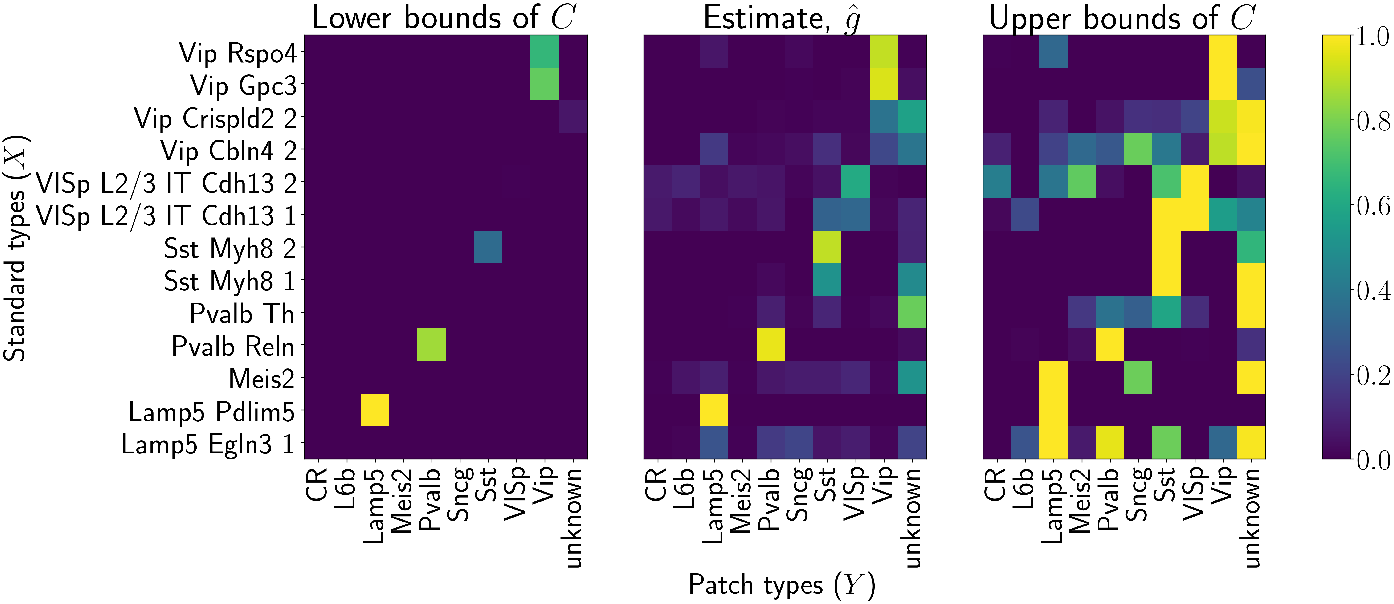
MLM estimation of the link, with credible interval. The central plot shows a portion of the MLM estimator applied to the data described in Figure 3. We examine a subselection of Standard types (*X*) and all of the Patch types (*Y*). For each combination *x, y* we draw a rectangle whose color indicates the value of the estimator *ĝ(y|x)*. We also determine the MLM’s confidence about these estimators. On the left we indicate the lower bounds indicated by the credible interval; on the right we indicate upper bounds. For some aspects of the link the intervals are much tighter than others. For example, it appears we have high confidence the Standard type ‘Vip Rspo4’ is highly associated with the Patch type ‘Vip.’ In contrast, we have almost no idea what is associated with the Standard type ‘Meis2.’ The upper credible interval bounds suggest that *g*(*y*|Meis2) could be nearly 1 for many different Patch types. Obviously it cannot be 1 for all of those types simultaneously, since Σ_*y*_ *g*(*y*|Meis2) = 1, but the data simply doesn’t tell us which *y* carries the mass.

The variability in the credible region suggests how to more closely determine the value of the link. For example, the significant ambiguity for cells with standard type ‘Lamp5 Egln 1’ suggests we need more distinct selection procedures that include these cells. If we could find a selection procedure that obtained many cells which measure as Standard type ‘Lamp5 Egln 1’ but no cells which measure as Patch type ‘Pvalb,’ this would show that the Standard ‘Lamp5 Egln 1’ is not associated with Patch type ‘Pvalb.’ On the other hand, if we could find a selection procedure that obtains many ‘Lamp5 Egln 1’ cells but *only* cells with Patch type ‘Pvalb,’ this would show the opposite. Once such additional selection procedures have been determined and experiments run, the MLM can be applied to the new data to determine what aspects of the link are still ambiguous.

## 4 Relation to prior work

The MLM infers the link between different measurement procedures to combine multimodal experimental data. There is a long line of literature on this subject. For example, when experiments are performed in batches, the exact measurement procedures can vary slightly between batches. The entire field of ‘batch effects’ is devoted to handling these problems. The general approach is to use some knowledge of the procedures to make modeling assumptions about the links. These assumptions give us a way to estimate the link (cf. [11]). If different measurement procedures yield results in the same space, we can also implicitly articulate these kinds of assumptions by assuming measurement procedures should yield results that are somehow ‘close.’ This leads to optimal transport techniques that use a distance measure to produce a link (cf. [12]). From the most general point of view, we are engaged in meta-analysis; we refer the reader to [13] for a general introduction to the field. The main distinguishing characteristics of this paper are two-fold: we place no assumptions on the nature of the link and focus on the resulting identifiability issues [14].

Our fundamental approach takes its origins from the causality literature. The MLM treats measurements never performed as latent random variables; this is a common approach in the causality community, and generally goes by the name of ‘Potential Outcomes’ (cf. [15]). The idea of using selection procedures also has precedent in the causal literature; it is sometimes referred to as ‘stratification by covariates’ (cf. [16]). The Markov Link Method can be understood as an application of these ideas to cases where the MLM assumption holds.

The main technical contribution of this paper is a method to translate the MLM assumption into practically useful credible intervals. In this we were inspired by a large literature of examples where assumptions are used to bound potentially unidentifiable parameters. Some of this literature also comes from the field of causality. For example, in [17] Bonet produces regions not unlike the ones seen here to explore whether a variable can be used as an instrument. The Clauser-Horne-Shimony-Holt inequality was designed to help answer causality questions in quantum physics, but it also sheds light on what distributions are consistent with certain assumptions [18]. More generally, the physics literature has contributed many key assumptions that bound unidentifiable parameters (cf. [19], [20], and the references therein). The closest work to this one would be [21], which uses two marginal distributions to get bounds on a property of the joint distribution (namely the distribution of the sum). We advance this approach to a more general-purpose technique, both by using many subpopulations to closely refine the MLM estimates and by considering the entire space of possible joint distributions instead of a single property of the joint.

## 5 Conclusion and future work

In this work, we formalize the concept of a ‘measurement link’ between two different types of experimental data. We develop the Markov Link Method (MLM), a tool to estimate this link. Critically, the MLM does *not* require data where both measurement procedures are applied to the same specimen. Thus the MLM can be applied even when measurement techniques are destructive, or in cases where obtaining multiple measurements from the same specimen is prohibitively costly.

To accomplish this, the MLM requires a variety of *selection* procedures; these selection procedures choose data from different (though perhaps overlapping) subpopulations and therefore provide different views into how the measurement procedures are related. The MLM combines many such views to optimally constrain the measurement link.

In this work the MLM is used for measurements that produce one of a finite set of values, such as procedures which measure a cell to determine its cell type. It is conceptually straightforward to extend the MLM to other kinds of measurement procedures, such as those that produce real values. Similarly, we demonstrated that the MLM can estimate the link between two measurement procedures. It is also straightforward to extend this approach to more than two types of measurements. We describe these extensions in Appendix D, but note that significant future effort maybe required to put these concepts into practice.

In summary, the MLM provides a generic tool to combine data across different experimental modalities. Every scientific experiment provides a glimpse into the domain under study. Tools that can combine these perspectives, such as the MLM, are critical to using all of our data to form accurate and coherent scientific theories.

## Acknowledgments

J.L. and L.P were supported by the following grants: Chan Zuckerberg Initiative 2018-183188, ONR N00014-17-1-2843, NIH U19 NS107613-01, NSF NeuroNex DBI-1707398, and the Gatsby Charitable Foundation. T.B., U.S., G.M., and H.Z. were supported by the Allen Institute for Brain Science. D.B. is supported by ONR N00014-11-1-0651, DARPA PPAML FA8750-14-2-0009, the Alfred P Sloan Foundation, and the John Simon Guggenheim Foundation.

## A Exact details of the Markov link method

Consider experiments yielding an Ω*_ℓ_* × Ω*_X_* matrix 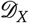 and an Ω*_ℓ_* × Ω*_Y_* matrix 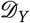, carrying the distribution

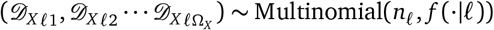

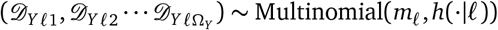

where *f*(*x*|ℓ), *h(y*|ℓ) are conditional distributions and there is some *g*(*y|x*) such that *h(y*|ℓ) = Σ*_x_f*(*x*|ℓ)*g*(*y|x*). This is simply a restatement of the assumptions we have made throughout this paper about how the objects of interest (*f, g, h*) are related to the data we can observe 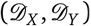.

The purpose of the MLM is to make estimates about *g* using the data 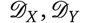. Unfortunately, *g* cannot be directly determined from the data. Even perfect knowledge of *f, h* may be insufficient to determine the true value of *g*. Some examples are detailed in Appendix C. This problem is called ‘nonidentifiability,’ and it can have some troubling consequences. For example, standard Bayesian analyses applied to nonidentifiable parameters will be extremely sensitive to the precise choice of prior beliefs. Even with infinite data, the prior beliefs may have a significant impact on inferences. To avoid these difficulties, we focus on objects that we know we can identify from data. In particular, we will look at lower bounds, upper bounds, and something in-between.

Let Θ(*f, h*) = {*g*: *h(y*|ℓ) = Σ*_x_f*(*x*|ℓ)*g*(*y*|*x*)} denote the set of links which are consistent with *f, h* and the Markov link method assumption. We define

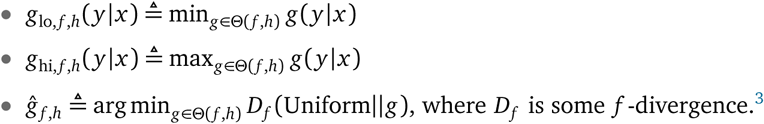

Even if *g* is nonidentifiable, *g*_lo_, *g*_hi_ may still be identifiable, and these quantities bound the measurement link according to the inequalities *q*_lo_(*y*|*x*) ≤ *g*(*y*|*x*) ≤ *g*_hi_(*y*|*x*). The estimator *ĝ* is also identifiable and we can also hope it will strike a middle ground. In producing this single point estimate we had to decide how to deal with the fundamental fact that actually any *ĝ* ∈ *Θ* might be correct. At a basic level, we could make two kinds of mistakes. We might claim a very strong association between the measurement procedures even though actually there is none. We might claim a very weak association even though actually there is a strong association. We choose to err on the side of asserting weak associations, by choosing the *g* which is as close as possible to uniform. We made this choice in the spirit of the Maximum Entropy Principle, i.e. that in the absence of other information we assume *X* is associated with each *Y* equally. This is perhaps as reasonable as any way to pick a particular *ĝ*. However, we reiterate that *ĝ* is just one possibility among many. It is safest to consider the full spectrum of possibilities by looking at the extremes *g*_lo_, *g*_hi_.

If we had perfect knowledge of *f, h*, the objects *g*_lo,f,h_, *g_hi,f,h_, ĝ_f,h_* would give us a reasonable understanding of what we can know about the link *g*. However, in practice we do not have access to *f, h*. Instead, we have access to the data 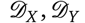 which enables us to estimate *f, h*. To account for uncertainty about these estimates, we take a Bayesian perspective. For prior beliefs about *f, h*, we take a noninformative uniform prior 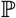:

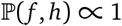

Following the Bayesian philosophy, we then incorporate new knowledge by conditioning. We have two important pieces of knowledge about *f, h*. First, we have observed the data, 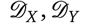. Second, we know from the MLM assumption that there exists *some* value *g* such that Equation (2) holds. We would like to condition on both of these facts. However, due to the Borel-Kolmogorov paradox, ‘conditioning on the MLM assumption’ is not a meaningful idea. Instead, it is necessary to define a variable indicating how much the MLM assumption fails, and condition on this variable being zero. In particular, let *D(h||h’)* denote the Kullback-Leibler divergence and Γ*(f,h)* = inf_*h’: Θ(h,h’)≠∅*_*D(h||h’)*. Let rf denote the event that Γ*(f, h)* = 0. Posterior uncertainty about *f, h* can then be articulated through the distribution

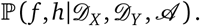

In terms of this posterior, we define the MLM point estimate *ĝ* and uncertainty bounds *C* as follows:

1. *ĝ* is calculated using posterior expectation:

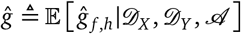

2. *C_x,y_* is calculated in terms of credible intervals. For each *x, y*, we define *C_x,y_* as the interval from the 2.5th percentile of *g*_lo,*f*,*h*_(*y*|*x*) to the 97.5th percentile of *g*_hi,*f,h*_(*y*|*x*) under the posterior distribution.

In practice, we were not able to find a way to compute these objects exactly. Given samples from the posterior distribution 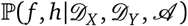, it would be straightforward to get good estimates. As seen in Appendix E, it is straightforward to compute *g*_lo_, *g*_hi_, *ĝ* from samples of *f, h*, so we could use use Monte Carlo approximations for our objects of interest. Unfortunately, it seems difficult to obtain samples from this posterior distribution. Common approaches to this type of problem involve Markov Chain Monte Carlo and Variational Inference, but we were unable to make these approaches work in practice. It seems nontrivial to work with the condition Γ*(p, h)* = 0 that formalizes the MLM assumption. We instead take a somewhat naïve approach. We start by drawing samples according to

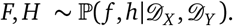

This can be achieved exactly, using the the conjugacy between the prior and the Multinomial distribution. Notice that these samples do not incorporate knowledge of the MLM assumption, insofar as they are not conditioned on the event Γ*(f, h)* = 0. To approximately remedy this, we define 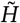 as the solution of min_h’_ *D(H|h’)*, subject to the constraint that Γ*(F, h’)* = 0. Optimization details can be found in Appendix E. We use the pair *F*, 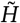 as *approximate* samples for the distribution 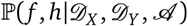. We can repeat this process to produce many samples of 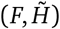 and use those samples to produce approximate Monte Carlo estimates for *ĝ*(*y|x*), *C_x,y_*. In the limit of large sample sizes we expect that *F, H* will nearly satisfy the MLM assumption in any case, so this approximation should not make a large difference. For example, on the transcriptomic dataset examined in the main text we found that the total variation distance between *H*(·|ℓ) and 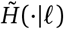 was about 15% (averaging over all selection procedures ℓ and various samples of *H*). For comparison, this is about three times smaller than the average total variation distance between *H*(·|ℓ) and *H*(·|ℓ’) for a randomly selected pair of selection procedures (ℓ,*ℓ’)*, which averages out to around 50%.

A summary of the final algorithm can be found below, in Algorithm 1. Consistency results for this final algorithm can be found in Appendix B.

**Algorithm 1:**
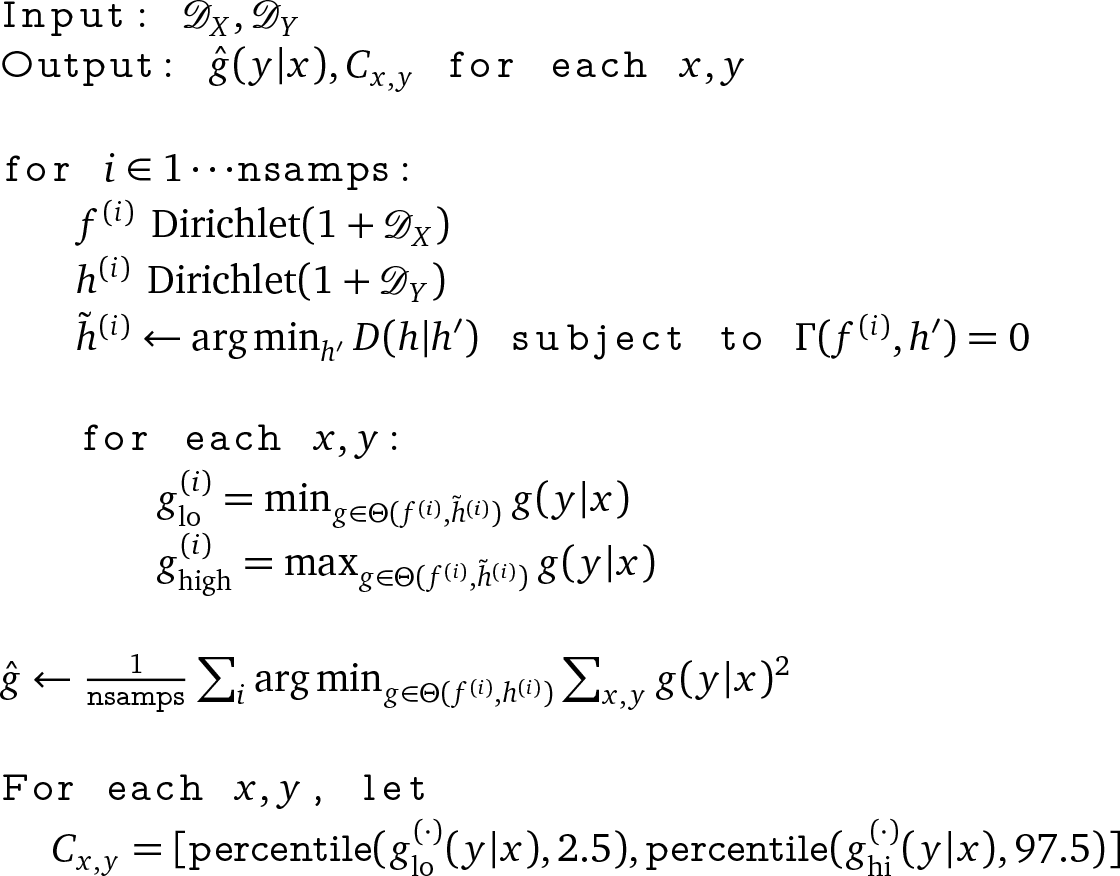
The Markov Link Method

Note that this algorithm includes solving several optimization problems. Details for how we solve those problems can be found below, in Appendix E.

## B Consistency results

We here show some fairly mild conditions under which the credible intervals *C_x,y_* defined in Algorithm 1 are asymptotically consistent, in that they are guaranteed to contain the true link parameters up to an arbitrarily small constant. Throughout this section we will adopt the notation found in that Algorithm. Our main assumptions are as follows:

- Linear independence. The matrix *B* defined by 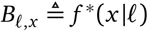 has linearly independent rows. This will typically occur whenever the number of selection procedures is no greater than the number of discrete values that the *X* measurement can return. Our use case featured a relatively small number of selection procedures, so we focus on that case.
- Positivity. The distribution *g*\* is strictly positive. This assumption greatly simplifies the theoretical analysis by allowing us to assume that many important objects are asymptotically normal.

We expect that these conditions are actually not necessary for our result. However they greatly simplify the analysis, yielding the following short consistency proof. In future work we hope to remove these conditions.

**Theorem 1**. By taking *n_ℓ_, m*_ℓ_ > *c* for *c* sufficiently large we can ensure *g*\*(*y|x*) ∈ *C_x,y_* ± *∈* with aribtrarily high probability and arbitrarily small *ε*> 0.

*Proof*. We first recall some classical results on posterior concentration for Multinomial data. Let *π* denote the posterior distribution on *f, h* under a uniform prior:

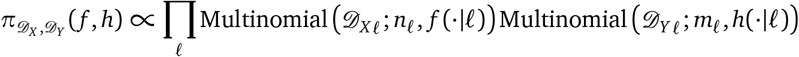

This is the key distribution used in Algorithm 1. Notice that if we consider 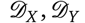 to drawn from multinomials parameterized by *f*\*, *h*\*, then 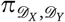 becomes a random measure. The randomness comes from the fact that 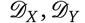 are considered to be random variables. In this setup, it is well-known that by taking *n, m* sufficiently high we can ensure that

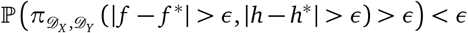

for arbitrararily small *∈*. Note that the exact norm chosen to define |*f* – *f*\*| is not particularly important; since these objects are finite-dimensional, all these norms are equivalent (e.g. *ℒ*^2^, total variation, uniform norm).

In order to account for our knowledge that there is some *g* such that Σ*_x_f*(*ℓ*|*x*)*g*(*y*|*x*) = *h*(*y*|*ℓ*), we do not work directly with the distribution π. Instead, recall that we define 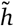 as as a solution of

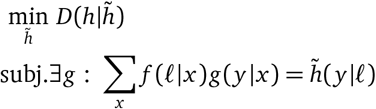

It is easily seen that this problem is strictly convex and so the solution is unique; thus 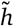 is a deterministic function of *f, h*. We would like to obtain a similar posterior concentration result for this altered variable, i.e. we can ensure 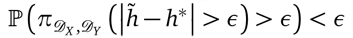 by taking *n*_ℓ_, *m*_ℓ_ sufficiently high. This follows because we already know *h*\* ≈ *h* and we can ensure 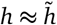 whenever *h* ≈ *h*, f* ≈ *h\**, using the positivity and linear independence assumptions. Indeed, we have that Σ*_x_f*\*(ℓ|*x*)*g*\*(*y*|*x*) = *h\**(*y*|ℓ), *f* ≈ *f* *, and *h* ≈ *h\**. We can therefore apply a kind of implicit function theorem result to show that we can find 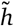 which is close to *h* in *ℒ*^2^ and such that ȃ*g* with 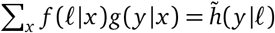 (note that the positivity condition ensures that this *ℒ*^2^ closeness is locally equivalent to the KL divergence which we actually minimize to find 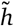). To apply such an implicit function theorem, we need to ensure two things: that the relevant Jacobians are invertible and that *h*\* does not lie on the boundary of the feasible space. In this case these conditions can be ensured by the independence of the rows of 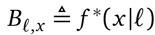 and the positivity of *g*, respectively.

We would now like to use the fact that *f* ≈ *f*\* and 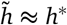 with high probability to show that *g*\*(*y|x*) ∈ *C_x,y_* ± *ε* with high probability. Without loss of generality, we will focus on the lower bound of the interval *C_x,y_*. Recall that this lower bound is defined as a percentile of the distribution of 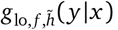.; Thus, it suffices to show that there is a high probability that 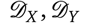 are such that 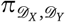 assigns high probability to 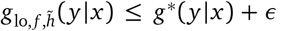. In light of the posterior concentration results above, it suffices to show that by taking 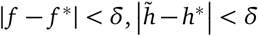 for *δ* sufficiently small we can ensure *g*_lo,f,h_(*y*|*x*) ≤ *g*\*(*y*|*x*) + *∈* for arbitrarily small *∈*. This is easily seen. Let 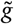 denote the *ℒ*^2^ projection to 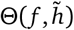. Once again the independence of the rows of *B* and the positivity allows us to ensure that 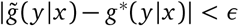 for every *x, y*. Thus, since 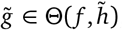, the very definition of *g*_lo_ yields that 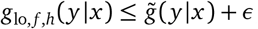, as desired.

## C Identifiability

The issue of identifiability comes up repeatedly throughout this paper. Here we give a brief overview of the fundamentals of this issue. We also present two suggestive case studies which we hope may inspire future research. In both cases we are able to prove something of interest - but not quite as much as we might hope. Here we will use the notation introduced in Appendix A.

First note that we can obtain arbitrarily good estimates of *f, h* by taking enough samples (i.e. taking *n*_ℓ_, *m*_ℓ_ sufficiently high). Let us therefore imagine for a moment that we in fact have *perfect knowledge* of *f, h*. Even so, the data do not necessarily tell us the value of the link *g*. There may be many possible links, *g*, which are all equally consistent with *f, h*. That is, we may have *g*_1_,*g*_2_ such that *h(y*|ℓ) = Σ_*x*_*f*(*x*|ℓ)*g*_1_(*y|x*) = *h(y*|ℓ) = Σ*_x_f(x*|ℓ)*g_2_(y|x)*. Both links yield the exact same distribution on the data we can observe, so there can be no way to use data to distinguish among them. This is known as a ‘nonidentifiability problem.’ Even with infinite data, we simply cannot identify exactly what the value of *g* might be.

We will now look at some examples:

### C.1 A simple failure case

Consider the case that Ω_ℓ_ = Ω*_Y_* = 2 and Ω*_X_* = 3. That is, there are 2 separate selection procedures, tool I recognizes 3 categories and tool II recognizes 2 categories. In particular, let us imagine that *f*(*x*|ℓ) = *A*_*ℓx*_ and *h*(*y*|ℓ) = *B*_*ℓy*_ where *A,B* are matrices given by

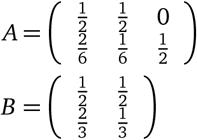

Rows correspond to different selection procedures and columns correspond to a different measurement outcome. Now let *g*(*y*|*x*) = *C_x,y_*, another matrix. The Markov link method assumption then tells us that *A* × *C* = *B*, where × indicates matrix multiplication. This corresponds to Ω_ℓ_ × Ω_*Y*_ = 4 equations. We also have a normalizing constraint that Σ*yg*(*y|x*) = 1, which creates Ω_*X*_ = 3 additional equations. However, these normalizing constraints actually make two of the MLM assumption constraints redundant. In the end, we have 5 constraining equations on the matrix *C*. However, the matrix *C* contains six numbers. The result is a degree of freedom in *C*, corresponding to an aspect of *g* that we simply cannot resolve. For example, here are two choices of *C* which are both consistent with the equation *A* × *C* = *B*:

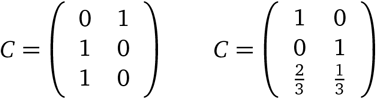

### C.2 Permutation matrices

Consider the case that Ω_ℓ_ = *k* and Ω_*X*_ = Ω_*Y*_ = 2^*k-1*^. That is, we are allowed to use *k* separate selection procedures, and measurement tools I and II can both return one of 2^k-1^ possible values. Let us furthermore assume that

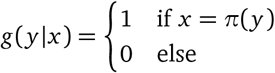

where *π* is a permutation on {1 … Ω_*Y*_}. Thus the two measurement procedures are deterministically related, but we don’t know which values of *X* correspond to which values of *Y*.

In this case, what selection procedures might we want to use to determine the permutation π? One natural idea would be to use a selection procedure that selects specimens taking on exactly *half* of the different values. We can easily imagine *k* such procedures, each selecting a different half of the values. The result is a set of selection procedures defined by *f* (*x*|ℓ) = *A*_*ℓ,x*_, where this matrix *A* is given by

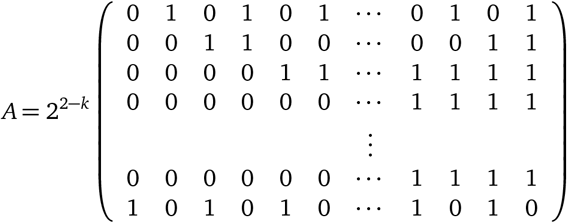

That is, the *x*th column of the first *k* – 1 rows is the binary expansion of the number *x* – 1, and the last row alternates 1s and 0s. Now let us say we have perfect knowledge of *f*(*x*|ℓ) and *h(y*|ℓ) = *Σ_χ_f* (*y*|ℓ) *g* (*y*|*x*)· Notice that due to the simple structure of *g* we obtain *h*(*y*|ℓ)= *A*_*ℓ,π(y)*_. However, let us imagine we know nothing about the true value of *g*.

How much can we say about *g*, if we only had knowledge of *f* and *h*? On the one hand, we observe that in the absence of any other constraints, the object *g* has 2^*2k–3*^ degrees of freedom. This is because there are 2^*k–1*^ values of ℓ and for each subpopulation *g*(·|ℓ) must lie in the 2^*k–2*^-dimensional simplex on 2^*k–1*^ atoms. On the other hand, we see that the Markov Link Assumption gives us *k* × (2^*k–1*^ – 1) linear constraints on the value of q. Indeed, for each subpopulation in 1 … *k* and each value of *y* ∈ 1 … 2^*k–1*^ we have an equation of the form

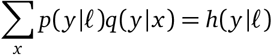

Of these k×2^*k*–1^ constraints, *k* of them are redundant with the fact that Σ*_y_g(y|x)* = 1. Thus, altogether, the Markov Link Assumption together with approximate knowledge of *p,h* gives us *k* × (2^*k*–1^ – 1) linear constraints. It would follow that *q* would have 2^2*k*–3^ – *k* × (2^*k*–1^ – 1) degrees of freedom yet remaining.

In conclusion, a simple degrees-of-freedom counting argument would suggest that there will be substantial ambiguity about what value *q* might take on, if our only knowledge about *q* is that it must satisfy Σ*_x_f*(*y*|ℓ)*q(y*|*x*) = *h(y*|ℓ). Indeed, we have *exponentially many* more degrees of freedom than we have constraints.

However, the reality is that *q* is exactly determined by *f, h*. This is possible because there are inequality constraints which also govern q, namely *g*(*y|x*) ≥ 0. Thus, while a simple degrees-of-freedom counting argument might suggest that we would have substantial identifiability issues in this problem, the reality is quite the opposite. This idea is made rigorous in the following theorem.

**Theorem 2**. Let *f, h* be as they are defined above. Then there is exactly one g that is consistent with *f, h* and the Markov Link assumption. That is, *g* is the only possible value satisfying

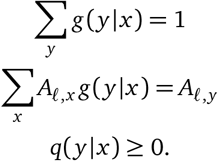

*Proof*. We prove by recursion. First take the case *k* = 2. In this case the result holds trivially, since *X, Y* ∈ {1}.

Now consider a general case *k* > 2. Without loss of generality, we take the simple case that π(*y*) = *y*, but the following arguments will hold for any π. Let us now focus on the constraints implied by the second-to-last row population. It is straightforward to see that these constraints imply

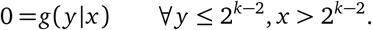

Indeed, for each *y* ≤ 2^*k*–2^ we obtain a constraint showing that Σ*_χ_>_2k–2_ q(y|x)* = 0, which yields that in fact *g(y|x)* = 0 for every *x* > 2^*k*–2^ and every *y* ≤ 2^*k*–2^.

It follows that for *y* ≤ 2^*k*–2^ the original constraints may be rewritten as

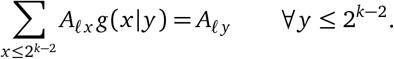

This is an example of the same problem we started with - except with *k* one smaller. Applying the inductive hypothesis, we may thus obtain that *g*(*y|x*) is uniquely determined for the first 2^*k*–2^ values of *x, y*. Moreover, since Σ_*y*≤2^*k*–2^_*g*(*y*|*x*) = 1, we see that *g* must also satisfy *g(y|x)* = 0 for *y* > 2^*k*–2^ and *x* ≤ 2^*k*–2^. Thus we have seen that *g* is uniquely identified for all entries except those in which *x, y* ≥ 2^*k*–2^.

For *x, y* ≥ 2^*k*–2^ we linearly combine equations concerning the first, last, and second to last rows of *A* with factors of 1,1,–1 respectively. We obtain constraints showing that Σ_*x*≤_2_^*k*–2^_*g*(*y*|*x*) = 0 for each *y* > 2^*k*–2^. We can then use the same reasoning to obtain that *g* is uniquely identified for the remaining values of *x, y*. □

This result is somewhat robust to slight perturbations in *f, h*. In particular, if we have some 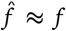 and 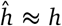 then at each stage of the argument we can replace statements of the form *g*(*y|x*) = 0 with statements of the form *g(y|x)* ≤ *ε*. Applying this with the kinds of arguments above will show that we can be sure that every point in 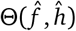 is arbitrarily close to *g* if we know that 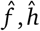 are sufficiently close to *f, h*.

However, it turns out that the relationship between *g* and *f, h* is not robust in every situation. In the next section we will see that it can in fact be quite discontinuous.

### C.3 Discontinuity

Consider the case that Ω_ℓ_ = 1 and Ω_*X*_ = Ω*_Y_* = 2. That is, there is only one selection procedure (no subpopulations) and both tool I and tool II can return one of 2 possible values. We will now consider two possiblities:

1. First let us take the case

- 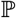(*X* = 1) = *f* (1) = 0
- 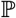(*X* = 2) = *f* (2) = 1
- 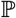(*Y* = 1) = *h*(1) = 0
- 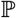(*Y* = 2) = *h*(2) = 1

In this case the MLM assumption Σ*_x_f*(*x*)*g*(*y*|*x*) = *h(y*) can be used to prove that *g*(1|2) = 0,*g*(2|2) = 1, but we now have *absolutely no* knowledge of *g*(1|1),*g*(2|1). This is because we simply never observed the case *X* = 1 (it occurs with probability zero), and so we cannot possibly have any knowledge about *g(y|x)* for *x* = 1.

2. Now let us take a slight variation:

- 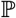(*X* = 1) = *f* (1) = 0.01
- 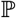(*X* = 2) = *f* (2) = 0.99
- 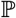(*Y* = 1) = *h*(1) = 0
- 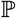(*Y* = 2) = *h*(2) = 1

In this case we can again prove that *g*(1|2) = 0, *g*(2|2) = 1, but we can also prove that *g*(1|1) = 0, *g*(2|1) = 1.

3. Now we take yet another slight variation:

- 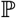(*X* = 1) = *f* (1) = 0.01
- 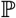(*X* = 2) = *f* (2) = 0.99
- 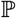(*Y* = 1) = *h*(1) = 0.01
- 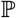(*Y* = 2) = *h*(2) = 0.99

In this case we can prove that *g*(1|2) ≤ 1/99 and *g*(2|1) ≥ 1 – 1/99, but we again cannot prove almost anything about *g*(2|1). In particular, it is easy to produce cases in which *g*(2|1) = 0 and other cases in which *g*(2|1) = 1.

The disturbing thing about this example is that by making infinitesimal perturbations to *f* we can pass from uncertainty to complete certainty back to uncertainty. It is for this reason that in this paper we refuse to ever treat *f, h* as fixed and given, always considering the space of perturbations around any such values.

It is worth noting that these kinds of problems essentially vanish if the true *g* is bounded away from zero i.e. *g(y|x)* > *c* for every *x, y* for some *c* > 0. This observation is the basis for our consistency result in Theorem 1 in Appendix B.

## D Extensions

In this paper we focused on the case of only two measurement procedures. We furthermore assumed that the measurement procedures could only return one of a finite number of results; in particular, we focused on the case that the procedures determined a ‘cell-type’ among a finite set of types. In this appendix we point the way to applying the ideas in this paper to more general problems.

### D.1 Setup

In general, the MLM begins with data from a collection of experiments. As described in the paper, we assume that each experiment can be characterized by two components: a selection procedure and a set of measurement procedures. Throughout the entire collection of experiments, we will assume that there are

- Ω_ℓ_ distinct selection procedures.
- *M* distinct measurement procedures.

Abstractly, we can consider all of the experiments together as a single dataset. For every specimen *i* gathered in any of the experiments, we are interested in

1. *ℓ_i_*, the sampling strategy used to gather specimen *i*. For example, if *ℓ_i_* = 3 that would indicate that specimen *i* was gathered in an experiment that used the third sampling strategy.
2. *X_i1_*, the measurement that would have been obtained from specimen *i* if we had observed it with the first measurement procedure.
3. *X_i2_*, the measurement that would have been obtained from specimen *i* if we had observed it with the second measurement procedure.
4. ⋮
5. *X_iM_*, the measurement that would have been obtained from specimen *i* if we had observed it with the *M*th measurement procedure.

Although we are interested in all of these values, not all of them may be observable. In particular, we may not measure all specimens with all measurement procedures. For example, consider the case that several of the measurement procedures destroy the specimen in the process of measuring it; in this case it is impossible to measure a specimen with all of the different measurement procedures. From this point of view, we can think of ℓ, *X* as a dataset with missing data: *X_ij_* is unobserved if specimen *i* was not measured with measurement procedure *j*. This perspective is sometimes referred to with the term ‘potential outcomes.’[15] That is, in practice we must pick a small set of measurement procedures to actually perform, but we can nonetheless think about the potential outcomes we might have obtained if we had used different procedures.

We assume that each *X_i_* is independent, and governed by some selection procedure dependent distribution,

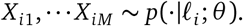

We will also assume that we have some prior on the unknown parameters of this distribution, *θ* ∼ *p(θ)*. In this paper we focused on the case that *p* was a categorical distribution which was parameterized in terms of *θ* = *f, g*. We placed a uniform prior on these unknown parameters. In general, we want to be able to consider any kind of distribution *p* for *θ, X*|ℓ.

#### D.2 Goal

In this more general setup, our goal is to infer some property of the joint distribution of *X*. There are many such properties one could be interested in. For example, one might wish to calculate the covariance between the results of two measurement procedures. Or perhaps one might be interested in the probability that three measurement procedures give the same result.

All aspects of a joint distribution can be analyzed by using so-called ‘test-functions.’ First, a question about the distribution is mathematically articulated by specifying a function. We then use statistical methods to estimate the expected value of that function:

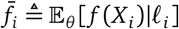

Our goal is to be able to estimate these kinds of expectations. Indeed, if we could determine *f̄_i_* for every function *f*, it is easy to show that we could use those estimates to determine the joint distribution governing *X* [22].

#### D.3 Challenges

If we had full knowledge of *X*, estimating *f̄* could be achieved with standard methods. However, we have ‘missing’ observations, because not every measurement procedure was performed on every specimen. The missingness of the data can cause *θ* to be unidentifiable. As emphasized in the main text, nonidentifiability can cause standard methods to fail. For example, one direct approach to estimating *f̄_i_* would be to apply Bayesian methods. We first compute the posterior distribution of *θ* conditioned on the data we can observe. We could then use this posterior to estimate *f̄_i_* by averaging over the posterior. However, conclusions from this posterior distribution can be extremely susceptible to the choice of prior. Even in the asymptotic limit of infinite data, non-identifiability causes the prior to have a continued impact on the conclusions. In high-dimensional settings it may be particularly difficult to reason about this prior, or determine which priors may or may not be sensible. For this reason, we advocate a different approach which is more robust to prior misspecification in the face of nonidentifiability.

#### D.4 Dealing with nonidentifiability

One solution is to take the nonidentifiability problem head-on. In particular, we define lower and upper bounds on our object of interest:

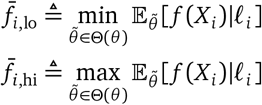

where θ(θ) indicates the equivalence class of parameters which yield the same distribution on the data we can observe. Note that we are guaranteed that *f*_*i*,lo_ ≤ *f̄_i_* ≤ *f*_*i*,hi_. Thus these quantities bound the true object of interest. These quantities are also identifiable, by definition. We can therefore apply traditional statistics to estimate these bounds. In particular, as in the main paper, we can construct credible intervals for these quantities using posterior samples of *θ*.

#### D.5 Introducing assumptions to tighten the bounds

For nontrivial problems, the lower and upper bounds introduced in the previous section may be extremely loose. They may offer very little insight into the true value of interest, *f̄*. This is the downside of taking this ‘head-on’ approach to identifiability. To tighten these bounds, we advocate introducing *hard constraints* that represent our beliefs and assumptions. We list some examples, below:

1. Distributional or smoothness assumptions. In this paper, every distribution was on a finite set, and we permitted these distributions to take the form of any categorical distribution. In applications involving continuous outputs, we may wish to assume particular distributions (e.g. a Gaussian assumption), or to place bounds on the smoothness of the output distributions.
2. The MLM assumption. The MLM assumption introduced in this paper can be generalized to the case of multiple measurement techniques. In general, it suffices to find a particular measurement modality which ‘statistically isolates’ the selection procedure from the other measurement modalities (without loss of generality, we will assume this is the measurement procedure corresponding to index 1). That is, we assume

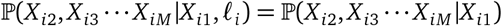

Conditional independence assumptions such as this can significantly tighten the bounds.

- Monotonicity assumptions. If measurement techniques 1 and 2 both measure essentially the same quality, we may wish to assume a stochastic monotonicity assumption. For example, we could assume that the distribution of *X_i2_*|*X_i1_* = *x* was first-order stochastically dominated by *X_i2_*|*X_i1_* = *x’* for any *x* < x’. Intuitively, this signifies that if *X_i1_* is bigger we expect *X_i2_* to be bigger, on average.

Applying these kinds of assumptions to real-world problems will not necessarily be trivial. It is difficult to predict which kinds of assumptions might yields bounds which are tight enough to be useful. In future work we hope to apply these ideas to a variety of datasets to make these ideas practical for general-purpose problems.

## E Numerical issues

There are three numerical problems which the MLM must solve. Here we detail our approach for solving each of these problems.

1. Projecting to the MLM assumption. Fix any values for *F, H*. One step in the MLM involves projecting *H* to the set of distributions which are consistent with *F* and the MLM assumption. In particular, we defined

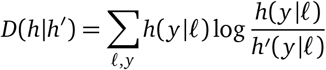

and we needed to solve the problem

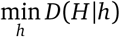

subject to the constraint that there exists some *q* such that *h*(*y*|ℓ) = Σ*_χ_F*(*x*|ℓ)*q*(*y|x*). Parametrizing valid *h* through *g*, we obtain the problem

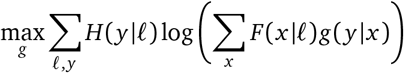

Taking derivatives one can readily show that this problem is convex. We solve it using exponentiated gradient ascent (cf. [23]). We initially guess that *g* is uniform. We then repeatedly make the updates

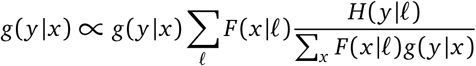

until convergence. The algorithm’s convergence criteria is that all parameters change less than 10^-5^ in a single iteration.

2. Linear programming. Fix any *f, h*. To deal with the identifiability issues, we defined 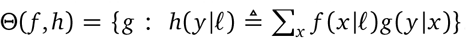. The MLM requires us to solve linear optimization problems within Θ, such as

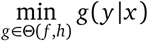

We solve these problems using the cvxopt python package.

3. Quadratic programming. To obtain the minimum *χ*^2^ divergence to uniform, the MLM also requires us to solve quadratic optimization problems within *Θ*:

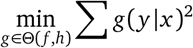

We solve these problems using the cvxopt python package

## Footnotes

1 Code to compute *ĝ*(*y|x*) and *C_x y_* is published at https://github.com/jacksonloper/markov-link-method, including a tutorial ipython notebook that details every computation made in this paper.

2 These are both technical assumptions which simplify the analysis. It seems likely that these conditions could be relaxed further, but we leave this for future work.

3 In practice, we choose a *χ*^2^ divergence because it makes the minimization problem a highly tractable quadratic program. See Appendix E for details.

